# Prolactin Modulates Renal Ischemia–Reperfusion Injury According to Sex and Cathepsin D Genotype

**DOI:** 10.64898/2026.09.09.750431

**Authors:** Maritza G. Verdugo-Molinares, Adriana Franco-Acevedo, Cesar O. Ramos-Garcia, Bibiana Moreno-Carranza, Norma Adan-Castro, Zesergio Melo

**Affiliations:** Biotecnología médica y farmacéutica, Centro de Investigación y Asistencia en Tecnología y Diseño del Estado de Jalisco, Guadalajara, México, 44270; Centro de Investigación Biomédica de Occidente, Centro Médico Nacional de Occidente, Instituto Mexicano del Seguro Social, Guadalajara, México; Division of Health Sciences, Universidad de Guadalajara, Centro Universitario de Tonalá, Tonalá, Jalisco; Monterrey Institute of Technology, Queretaro, Mexico; Neurobiology Institute, National University of Mexico (UNAM), Queretaro, Mexico; SECIHTI-Division of Health Sciences, Centro Universitario de Tonalá, Universidad de Guadalajara, Guadalajara, Mexico

**Keywords:** Renal ischemia-reperfusion injury, Prolactin, Cathepsin D, Sex differences, Kidney protection, Acute kidney injury

## Abstract

Renal ischemia–reperfusion injury (IRI) is a major driver of acute kidney injury and a critical determinant of long-term renal outcomes. Although sex differences in susceptibility to renal injury are well recognized, the hormonal mechanisms underlying these disparities remain incompletely understood. Prolactin (PRL), a pleiotropic hormone, has been implicated in both protective and deleterious renal effects, while Cathepsin D (CTD), a lysosomal protease, cleaves PRL into Vasoinhibins that possess antiangiogenic and immunomodulatory properties. The impact of CTD deficiency on PRL’s role in renal IRI has not been defined.

Here, we investigated whether CTD-regulated prolactin signaling modulates renal responses to IRI in a sex-dependent manner. Wild-type (WT) and heterozygous CTD mice of both sexes were subjected to unilateral renal IRI. PRL overexpression was achieved via lentiviral vector, and renal function, inflammatory cytokines, and angiogenic factors were assessed. In vitro Prolactin cleavage assays confirmed the enzymatic role of CTD. Proteomic analysis of renal cortex tissue was performed using label-free LC-MS/MS and pathway enrichment tools.

CTD cleaved PRL into Vasoinhibins at physiological pH, an effect blocked by Pepstatin A. PRL effects varied by sex and genotype. In WT females, PRL reduced renal dysfunction and increased IL-6 and TGF-β expression following IRI. CTD+/- females exhibited protection from injury, with PRL treatment reversing IL-6 induction. In males, PRL alone had modest effects, but CTD deficiency led to pronounced creatinine elevation, which PRL treatment normalized. Proteomic data revealed differential regulation of cytoskeletal and inflammatory proteins.

Cathepsin D heterozygosity modifies PRL’s biological effects on the injured kidney. The CTD–PRL axis regulates key inflammatory and angiogenic responses after IRI, revealing new mechanistic insights and potential therapeutic targets in acute kidney injury.

**Highlights:**

- Prolactin is cleaved into Vasoinhibins in the kidney.
- Prolactin exerts sex- and genotype-dependent protection in renal IRI models.
- VEGF and TGF-β expression is modulated by PRL in a Cathepsin D-dependent manner.
- PRL treatment reverses renal dysfunction in male mice.
- Proteomic and pathway analysis reveals inflammatory and metabolic modulation by PRL in the kidney.

## Introduction

Ischemia-reperfusion injury (IRI) is a major contributor to acute kidney injury (AKI) and remains strongly associated with progression to chronic kidney disease (CKD) and increased mortality. The pathophysiological cascade triggered by renal IRI involves profound cellular and structural alterations within the nephron, including endothelial dysfunction, oxidative stress, sterile inflammation, fibrotic remodeling, and activation of regulated cell death pathways [1, 2]. Despite the shared core features of IRI, the extent and progression of kidney damage can vary markedly depending on intrinsic host factors among which sex is increasingly recognized as a key biological variable [3, 4].

Epidemiological and experimental data consistently demonstrate that males are more susceptible than females to kidney injury and worse renal outcomes following IRI [4, 5]. This sex difference has been primarily attributed to the divergent effects of sex hormones on renal physiology [6]. In murine models, androgens have been shown to exacerbate kidney injury by promoting inflammation, fibrosis, and oxidative stress, while estrogens exert renoprotective effects through antioxidant, antiapoptotic, and vasodilatory mechanisms [7]. Indeed, Lima-Posada and colleagues have highlighted the protective role of estrogens in models of chronic kidney injury, reinforcing the relevance of sex-specific hormonal signaling in renal pathobiology [8]. However, beyond estrogens, the influence of other female-associated hormones such as prolactin (PRL) in the context of renal IRI remains largely unexplored [6].

PRL is a 23 kDa peptide hormone best known for its roles in lactation and reproductive biology [9], but it is also actively expressed in the kidney, particularly in epithelial and proximal tubular cells, where it participates in water and electrolyte handling [6, 10]. Elevated circulating prolactin levels are frequently observed in patients with chronic kidney disease and have been associated with adverse cardiovascular and renal outcomes [11]. The mechanistic implications remain unclear. Paradoxically, experimental evidence in non-renal models indicates that PRL may confer tissue protection. This hormone reduces infarct size and mortality in ischemic brain and heart injury, and promotes endothelial regeneration, vasodilation, and angiogenesis [12, 13]. Moreno-Carranza et al. demonstrated that PRL stimulates liver regeneration following partial hepatectomy in mice, an effect dependent on enhanced angiogenic signaling [14].

PRL’s vascular actions, however, are intricately regulated by its proteolytic fragments, collectively termed Vasoinhibins [13]. These bioactive peptides, ranging from 11 to 18 kDa, are generated through enzymatic cleavage of PRL by various proteases, with Cathepsin D (CTD), a lysosomal aspartyl protease, being the principal enzyme responsible [15]. Vasoinhibins exert potent antiangiogenic and proapoptotic effects [16]. These peptides inhibit vascular permeability, suppress nitric oxide production, and impair blood vessel formation [17]. Notably, Vasoinhibins have been detected in both serum and urine, and are known to accumulate in multiple tissues, including the kidney [18].

Cathepsin D is abundantly expressed in the renal cortex, particularly in glomeruli and along the tubular epithelium [19]. While its basal activity contributes to protein turnover and cellular homeostasis, excessive CTD activity has been implicated in a range of renal pathologies [20]. CTD-deficient mice exhibit early lethality and immune dysfunction, and podocyte-specific deletion of CTD leads to proteinuria, glomerulosclerosis, and podocyte apoptosis highlighting its essential role in glomerular integrity [19]. In pathological settings, CTD is upregulated in damaged kidneys and associated with fibrosis, apoptosis, and endothelial injury, particularly in patients with advanced CKD [20, 21].

Given the duality of prolactin signaling, mediated by both full-length hormone and Vasoinhibins fragments and the key role of CTD in their regulation, we hypothesized that Prolactin and Cathepsin D contribute to renal responses following IRI in a sex-dependent manner. In our study, we employ a murine model of renal IRI combined with genetic modulation of CTD and lentiviral PRL delivery to investigate how this hormonal and protease interaction shapes renal injury, inflammation, and repair.

## Methods

### Experimental Design and Animal Models

All experimental procedures were conducted in accordance with institutional ethical guidelines and were approved by the Institutional Committee for the Care and Use of Laboratory Animals (CICUAL) of the CIBO-IMSS (protocol number 1555). Male and female mice aged 12 to 15 weeks were used in this study. Two genotypes were studied: wild-type (WT) and heterozygous for Cathepsin D knockout (CTSD+/-, referred to as HT), on a C57BL/6 background (Figure 1b).

**Figure 1.**
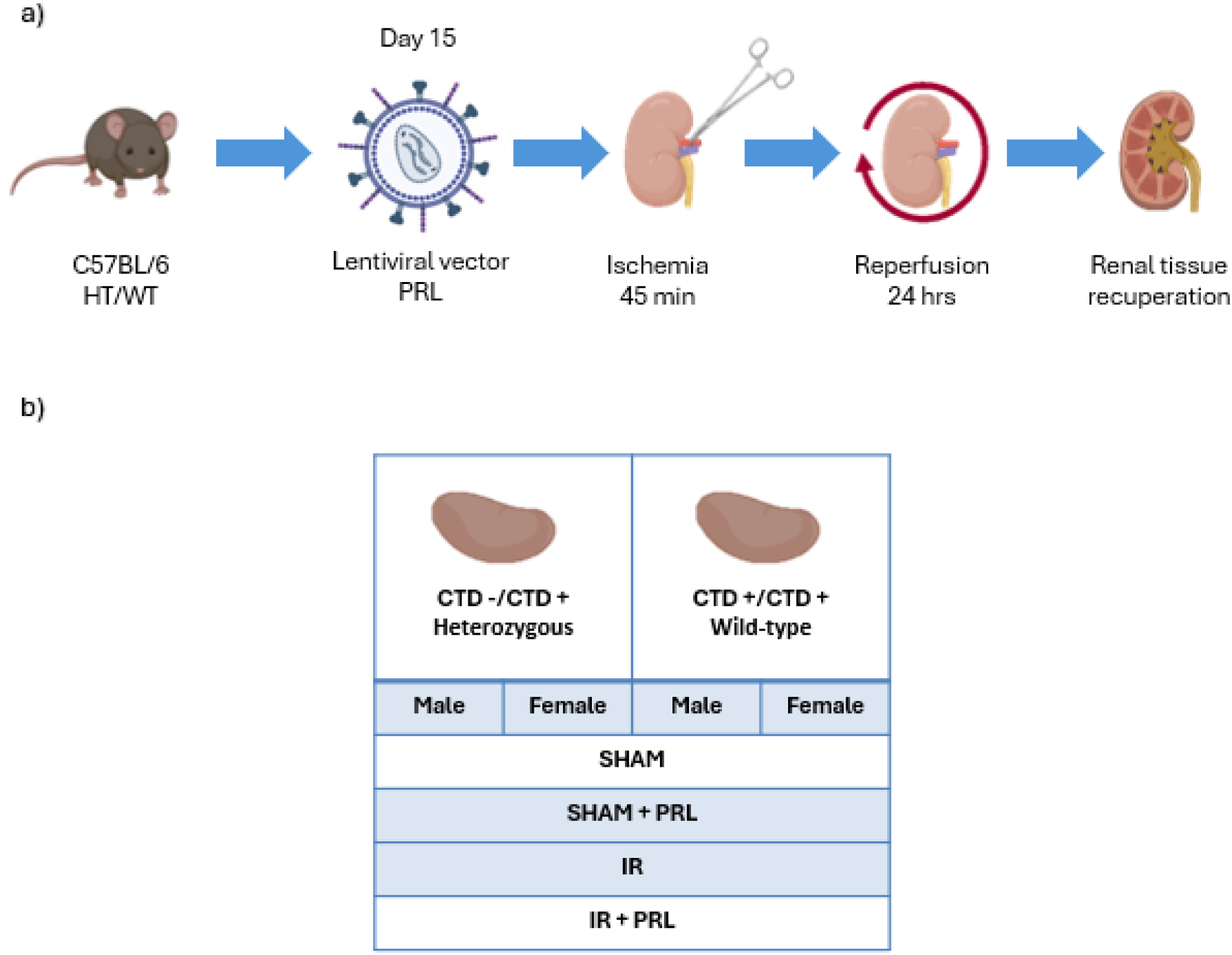
Experimental design and study groups. (a) Study workflow. Male and female C57BL/6 mice, wild-type (WT; Ctsd+/+) or heterozygous for Cathepsin D (HT; Ctsd+/-), received a single tail-vein injection of a lentiviral vector encoding mouse prolactin (LvPRL) or vehicle on day 15 before surgery. Unilateral left renal ischemia was induced for 45 min, followed by 24 h of reperfusion, after which renal tissue was collected. (b) Experimental groups. For each genotype and sex, animals were assigned to sham surgery, sham + PRL, ischemia-reperfusion (I-R), or I-R + PRL (n ≥ 5 per group). Renal tissue was analyzed for injury markers, inflammatory cytokines, angiogenic and fibrotic factors, and proteomic profiling. Schematic created with BioRender.com.

Mice were housed under standard laboratory conditions, 22a±n1°C, and 12 h light/dark cycle, with ad libitum access to food and water. Animals were randomly assigned to one of the following groups: sham-operated, IRI without treatment, or IRI with lentiviral prolactin overexpression (LvPRL). A minimum of 5 animals per group was used for molecular and functional analyses. Sample size was calculated based on prior power analyses for detecting differences in gene expression and creatinine levels. All efforts were made to minimize animal suffering and to reduce the number of animals used.

### Lentiviral Vector Construction and Delivery

To evaluate the effects of PRL overexpression on renal injury, a third-generation self-inactivating lentiviral vector encoding the mouse Prl gene was constructed under the control of the cytomegalovirus (CMV) promoter. The insert included the full-length murine Prl cDNA (NCBI Reference Sequence: NM_011164.4) cloned into the pLenti-CMV-GFP backbone, replacing the GFP cassette.

Lentiviral particles were produced by transient transfection of HEK293T cells using a three-plasmid system: the transfer plasmid (pLenti-CMV-Prl), the packaging plasmid (psPAX2), and the envelope plasmid (pMD2.G). Transfections were conducted using calcium phosphate precipitation, and viral supernatants were harvested at 48 and 72 hours post-transfection, filtered through 0.45 µm filters, and concentrated by ultracentrifugation at 25,000 rpm for 2 hours at 4°C. Viral titers were determined by p24 antigen quantification (QuickTiter™ Lentivirus Titer Kit, Cell Biolabs, Inc.).

Each animal received a single intravenous injection of lentiviral particles via the lateral tail vein at a dose of 1 x 10d transduction units in 100 µL of sterile PBS. The injection was performed 15 days prior to the induction of IRI, allowing sufficient time for stable PRL expression in target tissues. The timing and dose were established based on pilot experiments and previously validated protocols [22] to ensure robust systemic expression of PRL without inducing hyperprolactinemia-associated side effects. Successful expression of the transgene was confirmed by RT-qPCR and analysis of liver and kidney tissues collected at endpoint.

### Renal Ischemia-Reperfusion Injury Model

Unilateral left renal IRI was performed to evaluate the effects of PRL overexpression and Cathepsin D deficiency on acute kidney injury (Figure 1a). Mice were anesthetized with isoflurane at 4% and placed on a heated surgical table to maintain body temperature at 37D±D0.5D°C. Ophthalmic gel was applied to prevent corneal drying. A midline abdominal incision was made to expose left kidney. Ischemia was induced by clamping renal pedicles with atraumatic microvascular clamps for 45 minutes. The successful induction of ischemia was confirmed by a change in kidney color to dark purple. After the ischemic period, clamps were removed to allow reperfusion, which was visually confirmed by the return of normal coloration within 5 minutes. The abdominal wall and skin were closed in layers using absorbable 5-0 sutures. Sham-operated animals underwent the same surgical procedure, including exposure of the renal pedicles, but without application of the clamps.

To minimize postoperative pain, all animals received a single dose of buprenorphine 0.05 mg/kg, subcutaneously and were monitored until full recovery. After 24 hours of reperfusion, mice were euthanized by exsanguination under deep anesthesia. Blood and tissue samples were collected for biochemical and molecular analyses. Both kidneys were rapidly excised and rinsed in ice-cold PBS.

Tissues for protein extraction were homogenized in RIPA buffer (150 mM NaCl, 1% NP-40, 0.5% sodium deoxycholate, 0.1% SDS, 50 mM Tris-HCl, pH 7.4) supplemented with a protease inhibitor cocktail (Roche). Homogenates were incubated on ice for 30 minutes with periodic vortexing and then centrifuged at 14,000 × g for 20 minutes at 4°C. Supernatants were collected, quantified using the Bradford assay, and stored at –80°C for downstream Western blot or mass spectrometry. Total RNA was extracted using TRIzol reagent following the manufacturer’s instructions. RNA purity and integrity were verified by spectrophotometry. RNA samples were stored at –80°C for RT-qPCR analysis.

### Gene Expression Analysis (qPCR)

Total RNA was isolated from frozen kidney and liver tissues using TRIzol™ Reagent (Invitrogen) according to the manufacturer’s protocol. Briefly, 50–100 mg of tissue was homogenized in 1 mL of TRIzol, followed by chloroform extraction and isopropanol precipitation. The RNA pellet was washed with 75% ethanol, air-dried, and resuspended in RNase-free water. RNA concentration and purity were determined using an Epoch Biotek™ spectrophotometer. cDNA was synthesized from 1 µg of total RNA using the High-Capacity cDNA Reverse Transcription Kit (Applied Biosystems) in a 20 µL reaction volume, following the manufacturer’s instructions. Quantitative PCR was performed using SYBR™ Green PCR Master Mix (Thermo Fisher Scientific) on a StepOnePlus™ Real-Time PCR System (Applied Biosystems). Gene expression levels were normalized to the housekeeping gene Gapdh. Primer sequences were designed using online Primer-BLAST or obtained from validated references [23] and verified for efficiency and specificity via melt curve analysis. Relative expression levels were calculated using the 2^A^-AACt method.

Western blot assays were performed to evaluate prolactin cleavage and the presence of Vasoinhibins in vitro using rat renal extracts. Recombinant human prolactin (hPRL; 200 ng per reaction; R&D Systems) was incubated with 10 µg of total kidney protein extract for 24 hours at 37 °C in a reaction buffer at pH 7.0. Where indicated, Pepstatin A (10 µM; Sigma-Aldrich), a Cathepsin D inhibitor, was added.

After 24 h of incubation, samples were denatured in Laemmli buffer, resolved by SDS-PAGE on 15% polyacrylamide gels, and transferred to nitrocellulose membranes. Membranes were blocked with 5% non-fat milk in TBS-T (Tris-buffered saline, 0.1% Tween-20) for 1 hour at room temperature and incubated overnight at 4 °C with primary antibodies against Prolactin (1:1,000; Santa Cruz Biotechnology) and [3-Tubulin (1:5,000; Cell Signaling) as a loading control. After washing, membranes were incubated with HRP-conjugated secondary antibodies (1:10,000; Jackson ImmunoResearch) for 1 hour at room temperature. Signal detection was performed using enhanced chemiluminescence (ECL; Thermo Fisher), and bands were visualized on a ChemiDoc™ imaging system (Bio-Rad). Measurement of Serum Creatinine Blood samples were collected by cardiac puncture at the time of euthanasia and centrifuged to obtain serum as described above. Quantification was performed using a colorimetric enzymatic assay (Creatinine Assay Kit, BioAssay Systems) according to the manufacturer’s instructions. Briefly, 5 µL of serum was mixed with 200 µL of working reagent in a 96-well microplate, and the optical density (OD) was measured at 510 nm after 5 minutes of incubation at room temperature. A standard curve was generated using known concentrations of creatinine, and the concentration of each sample was calculated accordingly.

Assays were performed duplicate, and intra-assay coefficient of variation was below 5%. The detection range was 0.1 to 20 mg/dL. Results are expressed in mg/dL.

### Sample Preparation and Protein Extraction

Renal cortex tissue from each mouse was homogenized in RIPA lysis buffer (150 mM NaCl, 1% NP-40, 0.5% sodium deoxycholate, 0.1% SDS, 50 mM Tris-HCl, pH 7.4) supplemented with protease inhibitors. Homogenates were incubated on ice and centrifuged at 14,000 x g for 20 minutes at 4n°C. Supernatants were collected and total protein concentration was quantified using the Bradford method. 20 µg of protein per sample were subjected to trypsin digestion using the FASP (Filter-Aided Sample Preparation) protocol. Protein disulfide bonds were reduced with dithiothreitol (DTT) and alkylated with iodoacetamide (IAA). Samples were then passed through a 10 kDa filter unit (Millipore), washed with urea buffer, and digested overnight at 37°C with sequencing-grade modified trypsin (Promega) at a 1:50 enzyme-to-substrate ratio. Peptides were recovered by centrifugation, acidified with formic acid, desalted using C18 reverse-phase columns (Pierce™), and dried under vacuum.

### LC-MS/MS Analysis

Peptides were analyzed using a high-resolution mass spectrometer. Peptide separation was performed on an analytical column (25 cm × 75 µm, 1.6 µm C18 particles) using a linear gradient of 5%–35% acetonitrile with 0.1% formic acid over 120 minutes at a flow rate of 300 nL/min.

The timsTOF Pro was operated in data-independent acquisition (DIA) mode. MS data were acquired in PASEF mode (parallel accumulation–serial fragmentation) with ion mobility enabled. The mass range was set to m/z 100–1700, and MS/MS spectra were collected for the top 10 precursors per cycle. The databases were obtained from the Uniprot platform. Progenesis software was used to identify the signals.

### Pathway Enrichment and Network Analysis

Gene Ontology (GO) enrichment was performed using FunRich v3.1.3, focusing on biological processes, molecular functions, and cellular components. DEPs were uploaded as UniProt accession numbers, and enrichment was assessed against the mouse reference proteome. Pathways with p < 0.05 (adjusted by Benjamini-Hochberg FDR correction) were considered significantly enriched.

### Statistical Analysis

All quantitative data are presented as mean ± standard deviation (SD) unless otherwise indicated. For comparisons involving more than two groups, one-way ANOVA followed by Tukey’s post hoc test was used. When data did not meet normality assumptions, Kruskal– Wallis test with Dunn’s multiple comparisons was applied. For binary comparisons unpaired Student’s t test or Mann–Whitney U test was used as appropriate. Differences were considered statistically significant when p < 0.05. All statistical analyses were performed using GraphPad Prism v9.5.1 (GraphPad Software Inc.).

## Results

### Cathepsin D Expression Is Upregulated in the Kidney After Ischemia-Reperfusion Injury (IRI) in a Sex-Dependent Manner

Quantitative PCR analysis showed a significant increase in Cathepsin D mRNA levels following IRI in both male and female mice compared to sham controls. Notably, females exhibited a higher fold-change in CTD expression relative to males after IRI (p < 0.05) (Fig. 2a). This suggests a sex-specific response in the expression of CTD during kidney injury.

**Figure 2.**
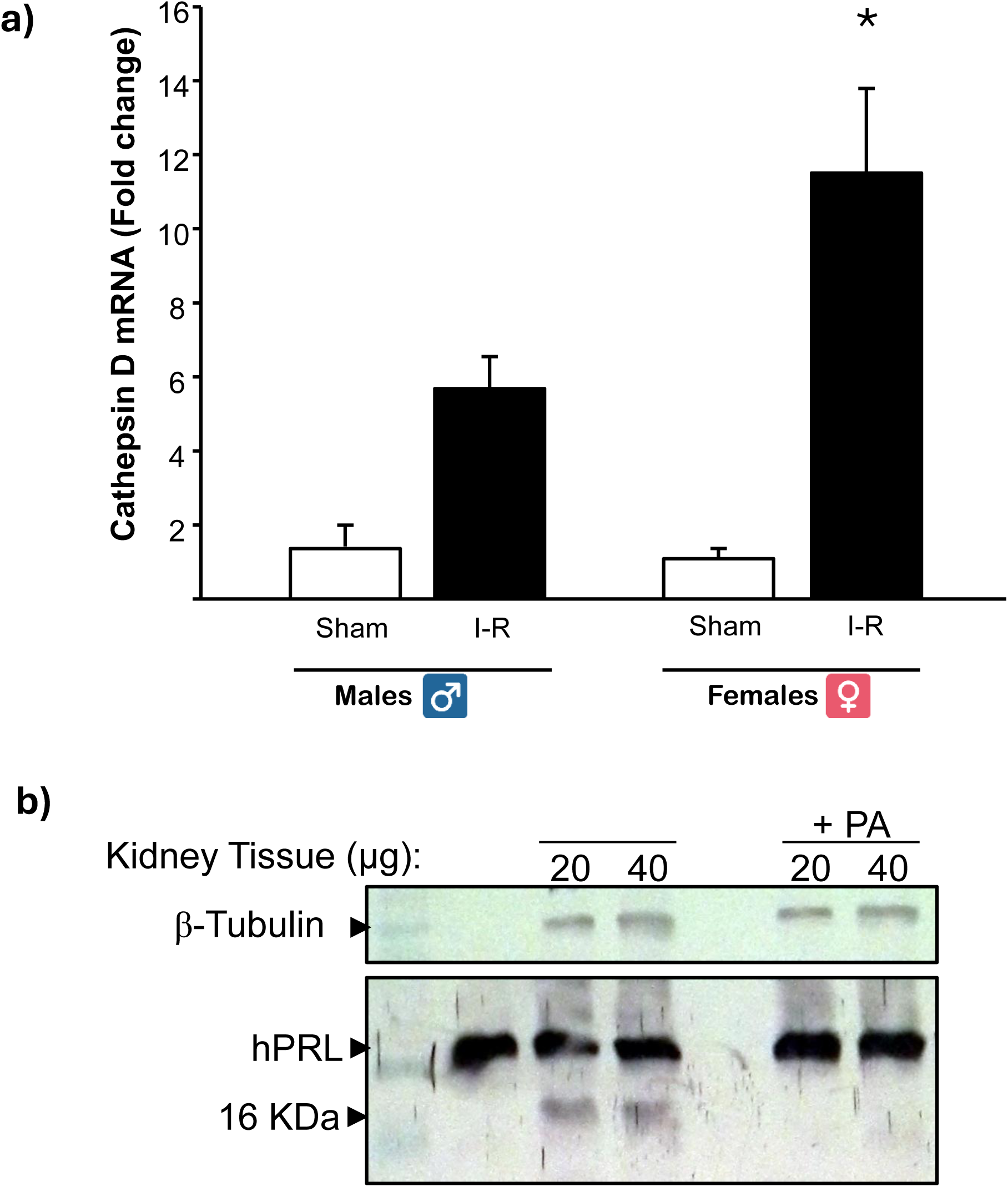
Cathepsin D is upregulated after ischemia–reperfusion injury and cleaves prolactin into Vasoinhibins. (a) Cathepsin D mRNA levels in the kidney of male and female mice after sham surgery or ischemia–reperfusion (I-R), measured by RT-qPCR and expressed as fold-change relative to sham (normalized to Gapdh; 2^A^-AACt). Cathepsin D expression increased after I-R in both sexes, reaching statistical significance in females (*p < 0.05 vs. the corresponding sham). Data are mean ± SD; n ≥ 5 per group. (b) In vitro prolactin cleavage assay. Recombinant human prolactin (hPRL, 200 ng) was incubated with kidney protein extract (20 or 40 µg) at pH 7.0 and 37 °C for 24 h, in the absence or presence of the Cathepsin D inhibitor Pepstatin A (PA). Immunoblotting shows conversion of full-length hPRL (∼23 kDa) into a ∼16 kDa Vasoinhibins fragment, which was prevented by PA; [3-Tubulin is shown as loading control.

### Cathepsin D Cleaves Prolactin into Vasoinhibins In Vitro

To assess whether CTD is responsible for PRL cleavage into Vasoinhibins fragments, hPRL was incubated at 37°C, pH 7.0, for 24 hours in the presence or absence of Pepstatin A (PA), a selective CTD inhibitor. Western blot analysis using kidney tissue protein (20–40 µg) showed that hPRL underwent cleavage, producing smaller molecular weight bands indicative of Vasoinhibins.

The addition of Pepstatin A inhibited this cleavage, preserving full-length PRL and reducing Vasoinhibins formation (Fig. 2b). This confirms that CTD enzymatically cleaves PRL under physiological conditions, and that this process can be inhibited pharmacologically.

### Prolactin Delivery via Lentiviral Vector Increases Liver Weight Relative to Body Mass

To evaluate potential systemic effects of PRL overexpression, we measured liver weight relative to total body weight in mice receiving either PRL-expressing or control lentiviral vectors. Mice treated with PRL lentivirus exhibited a modest but statistically significant increase in liver-to-body weight ratio compared to untreated controls (*p* < 0.05), suggesting that systemic PRL may influence hepatic physiology (Fig. 3a).

**Figure 3.**
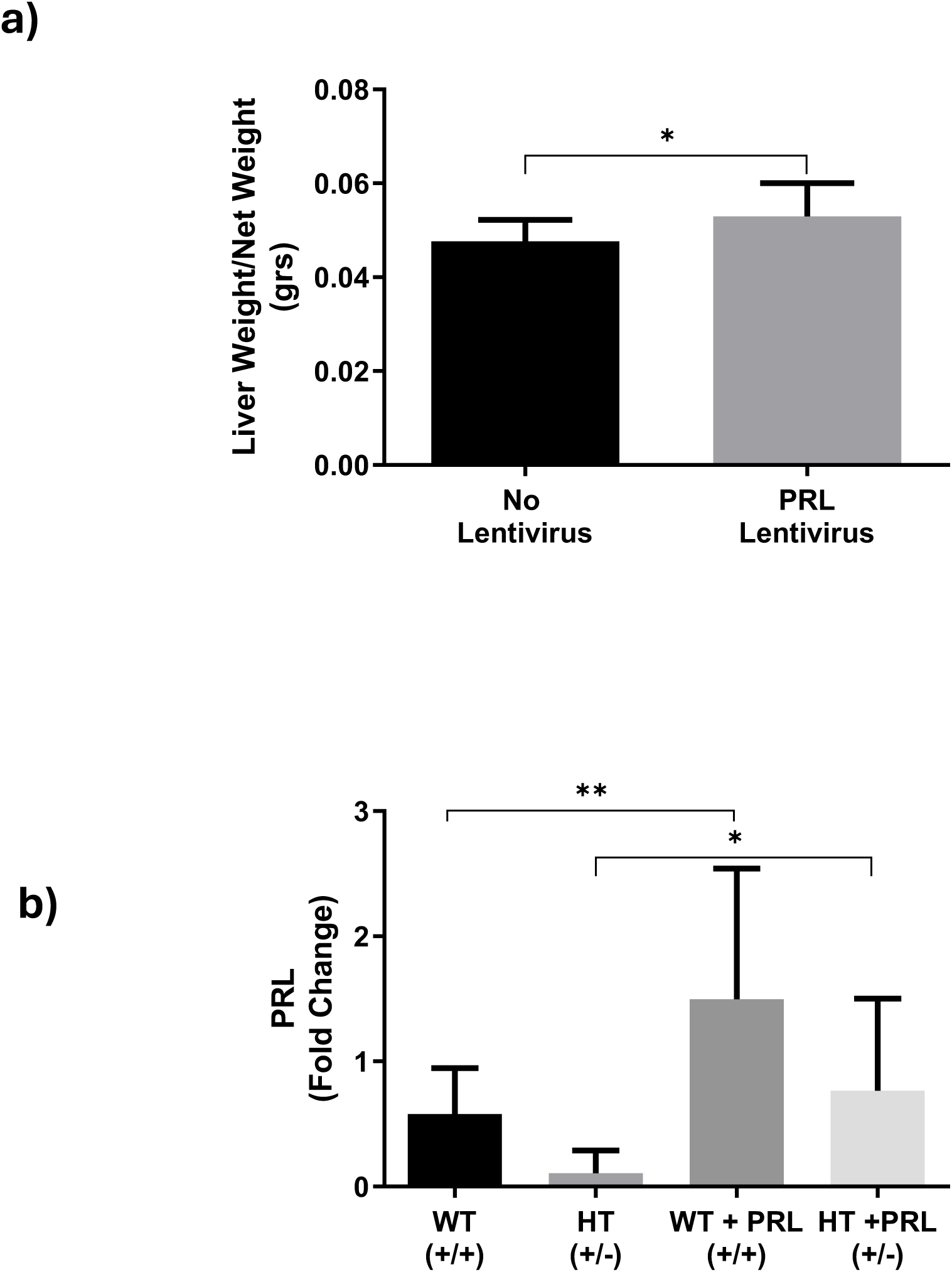
Validation of lentiviral prolactin overexpression and its systemic effect on liver mass. (a) Liver-to-body weight ratio in mice that received the vehicle or PRL-encoding lentivirus. PRL delivery produced a modest but significant increase in relative liver mass (*p < 0.05), indicating a systemic effect of the transgene. (b) Renal PRL mRNA levels (RT-qPCR; fold-change normalized to Gapdh) in wild-type (WT; Ctsd+/+) and heterozygous (HT; Ctsd+/-) mice, with or without PRL lentivirus. Baseline PRL expression was significantly lower in HT than in WT mice; lentiviral delivery increased PRL expression markedly in WT+PRL animals (**p < 0.01) and more modestly in HT+PRL animals (*p < 0.05). Data are mean ± SD; n ≥ 5 per group.

To confirm the efficacy of lentiviral-mediated PRL overexpression, PRL mRNA levels were quantified across WT and HT mice. In the absence of lentivirus, baseline PRL expression was significantly lower in HT mice compared to WT (**p** < 0.01). Lentiviral delivery of PRL resulted in a robust increase in PRL expression in WT mice (*p* < 0.05 vs. WT baseline), while PRL levels also increased in HT+PRL animals, though to a lesser extent. These results indicate that lentiviral delivery effectively upregulates PRL expression, with magnitude of induction partially influenced by genotype (Fig. 3b).

### Prolactin Modulates IRI-Induced Kidney Dysfunction in a Sex- and Genotype-Dependent Manner

Serum creatinine levels were significantly elevated in WT female mice following IRI compared to Sham WT controls (*p* < 0.001), indicating acute kidney dysfunction. Remarkably, treatment with PRL significantly reduced creatinine levels in I-R WT females (*p* < 0.01), suggesting a protective effect of PRL on renal function (Figure 4a).

**Figure 4.**
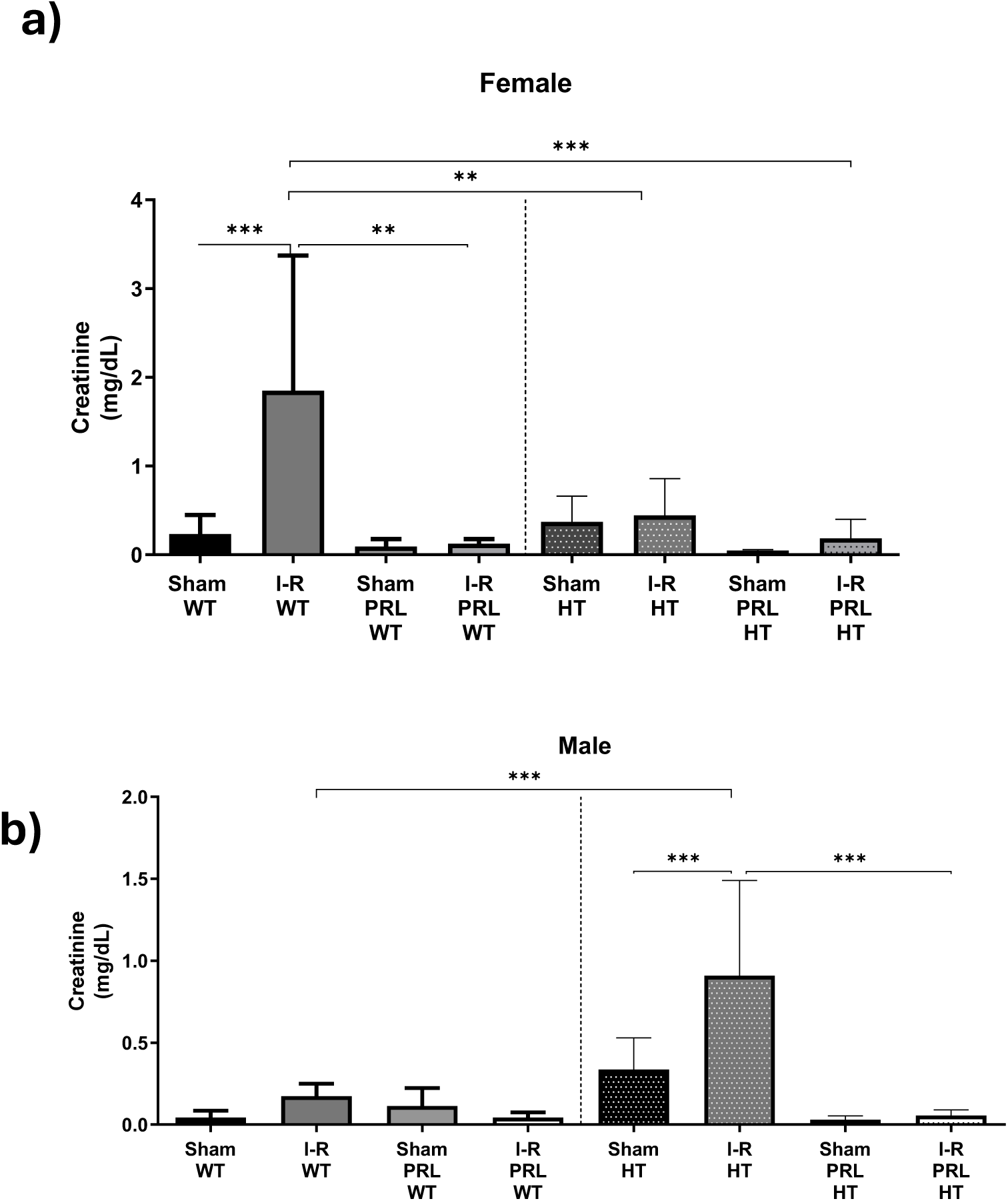
Prolactin modulates ischemia–reperfusion-induced renal dysfunction in a sex- and genotype-dependent manner. Serum creatinine (mg/dL) in (a) female and (b) male mice across genotype (WT, HT), injury (sham, I-R) and treatment (± PRL) conditions. (a) In females, I-R markedly increased creatinine in WT animals (***p < 0.001 vs. sham WT), an effect significantly attenuated by PRL (**p < 0.01); HT females were largely protected, showing minimal injury and no additional benefit from PRL. (b) In males, WT animals developed only mild dysfunction after I-R, whereas HT males showed a pronounced creatinine rise (***p < 0.001 vs. sham HT) that was fully normalized by PRL treatment (***p < 0.001). Data are mean ± SD; n ≥ 5 per group. *p < 0.05, **p < 0.01, ***p < 0.001.

In HT females, in which Cathepsin D-dependent PRL processing may be reduced, IRI did not significantly elevate creatinine levels compared to Sham, and the addition of PRL had no further impact. These findings suggest that PRL’s renoprotective effects in IRI are modulated by genetic background, probably involving Cathepsin D activity.

Overall, PRL treatment attenuated renal dysfunction in WT females subjected to IRI, supporting its protective role in female renal physiology (Figure 4a).

In WT male mice, IRI did not significantly elevate serum creatinine levels compared to sham controls, although a mild trend was observed. However, PRL treatment in the I-R WT group did not affect creatinine levels (Figure 4b).

In contrast, HT male mice exhibited a significant rise in serum creatinine following I-R (*p* < 0.001 vs. Sham-HT), indicating more severe kidney dysfunction. Importantly, PRL administration in I-R HT males significantly reduced serum creatinine to baseline levels (*p* < 0.001), fully preventing the IRI induced renal dysfunction.

These data suggest that PRL exerts a stronger renoprotective effect in HT males than in WT males under I-R conditions (Figure 4b), possibly due to altered regulation of CTD or Vasoinhibins activity in the HT model.

### Kidney Injury Molecule-1 (KIM-1) Expression Reflects Injury Severity and is Modulated by Prolactin and Genotype

KIM-1 expression was evaluated in male and female mice, genotypes and treatment conditions. In female mice (Figure 5a), KIM-1 levels were significantly elevated in the I-R WT group compared to Sham WT controls (*p* < 0.05), indicating injury. This induction was absent in the I-R Female HT group, suggesting that heterozygosity confers some degree of protection. Although no statistically significant differences were observed among PRL-treated groups, there was a trend toward reduced KIM-1 levels, particularly in the I-R Female HT+PRL group, which showed the lowest expression among the injury groups.

**Figure 5.**
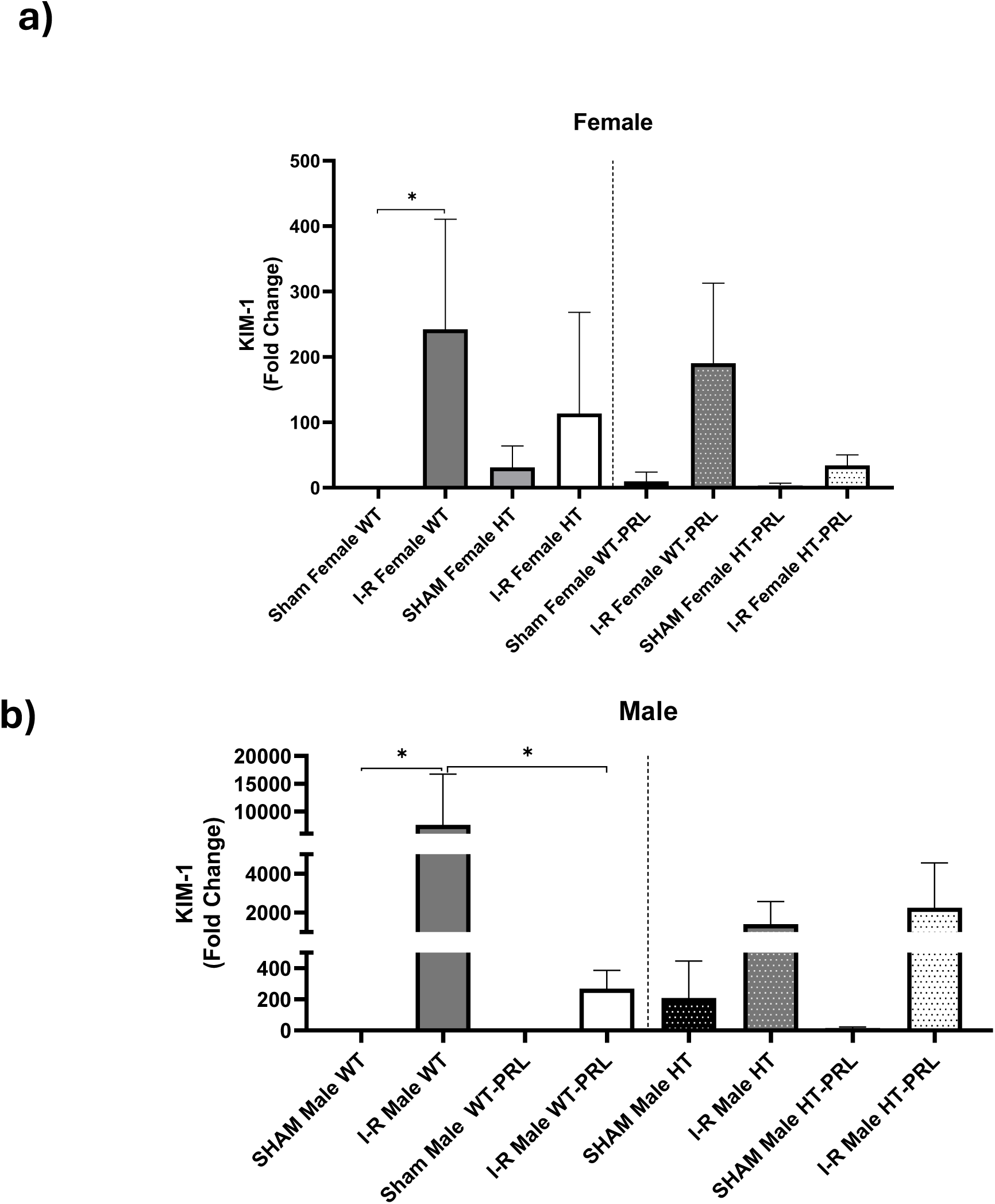
KIM-1 expression reflects injury severity and is modulated by prolactin and Cathepsin D genotype. Renal Kidney Injury Molecule-1 (KIM-1) mRNA (RT-qPCR; fold-change normalized to Gapdh) in (a) female and (b) male mice across genotype, injury and treatment groups. (a) In females, KIM-1 was significantly induced in I-R WT animals (*p < 0.05 vs. sham WT); this induction was absent in HT females, and PRL-treated groups trended toward lower expression. (b) In males, KIM-1 increased significantly in I-R WT animals relative to both sham WT and I-R WT+PRL (*p < 0.05), indicating attenuation by PRL, whereas HT males showed a uniformly blunted response. Data are mean ± SD; n ≥ 5 per group. *p < 0.05.

In male mice, KIM-1 expression increased significantly in I-R WT animals compared to both Sham WT and I-R WT+PRL groups (*p* < 0.05), suggesting that prolactin treatment attenuates injury severity. Heterozygous males (HT and HT+PRL) showed moderate and uniform expression levels, like sham controls, indicating an intrinsically blunted injury response (Figure 5b).

These results reveal a sexual dimorphism in KIM-1 expression following IRI and suggest that both PRL treatment and Cathepsin D genotype play protective roles, more prominently in males, but with discrete effects in females depending on genotype.

### HIF-1a Expression is Differentially Regulated by Genotype, Sex, and Prolactin Treatment

HIF-1a expression was assessed in all genotypes, sexes, and treatment groups to evaluate hypoxia responsive signaling after IRI.

In females, HIF-1a was basally elevated in Sham HT mice, displaying the highest fold change among all groups (Figure 6a). IRI modestly reduced this expression in HT animals, while PRL treatment uniformly suppressed HIF-1 expression in both WT and HT backgrounds regardless of injury status. WT females (both Sham and I-R) consistently exhibited negligible HIF-1a expression.

**Figure 6.**
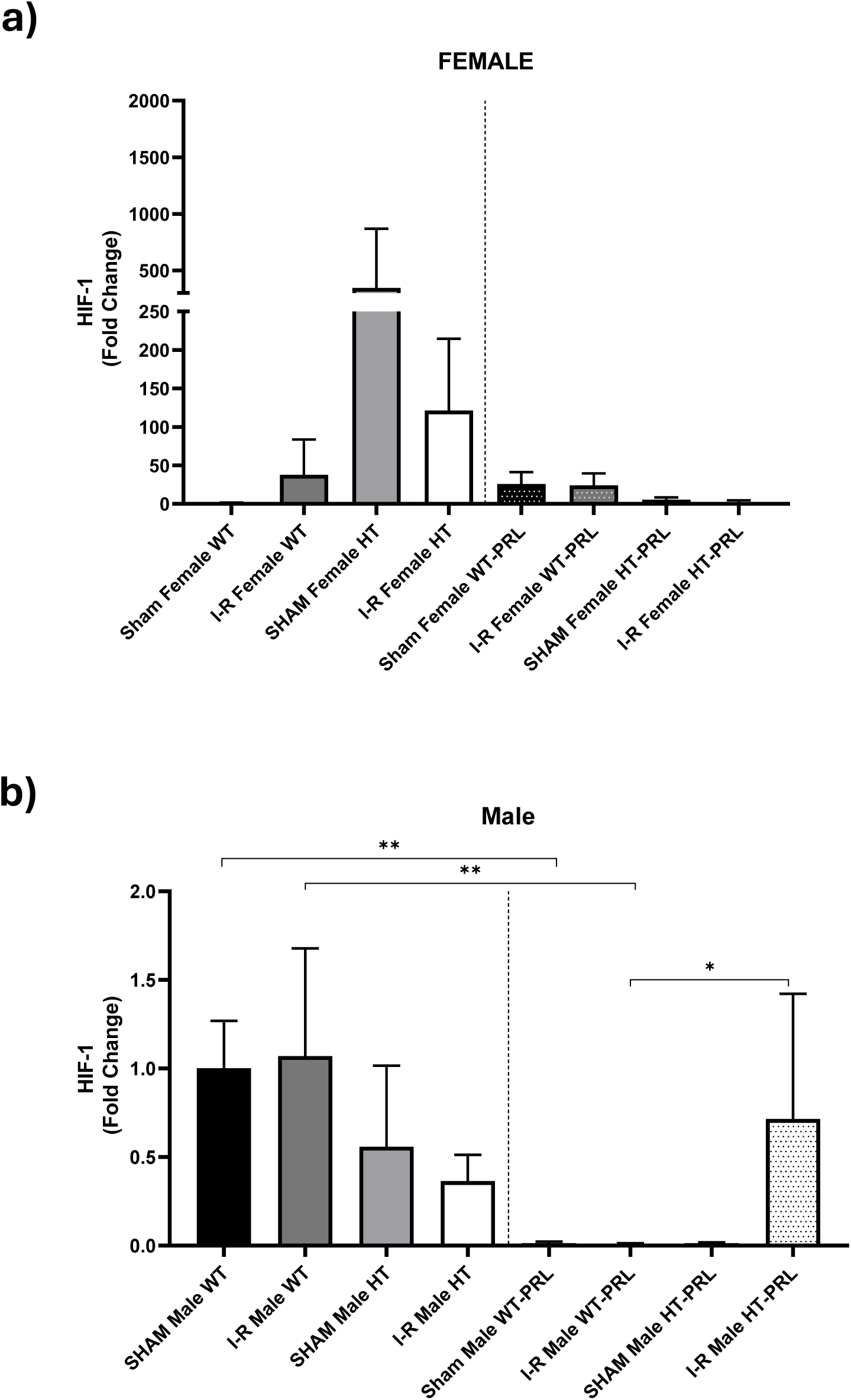
HIF-1a expression is regulated by genotype, sex and prolactin. Renal HIF-1a mRNA (RT-qPCR; fold-change normalized to Gapdh) in (a) female and (b) male mice. (a) In females, HIF-1a was basally elevated in sham HT animals and reduced by I-R; PRL treatment suppressed HIF-1a in both genotypes, whereas WT females showed consistently low expression. (b) In males, HIF-1a was significantly decreased in I-R HT relative to sham WT and sham HT (**p < 0.01); PRL restored HIF-1 in I-R HT males (*p < 0.05 vs. sham HT+PRL), while WT males treated with PRL showed minimal expression. Data are mean ± SD; n ≥ 5 per group. *p < 0.05, **p < 0.01.

In male mice, HIF-1a expression was significantly decreased in the I-R HT group compared to Sham WT and Sham HT controls (*p* < 0.01), suggesting that ischemia-reperfusion suppresses HIF-1a activation in heterozygous males (Figure 6b). Remarkably, PRL treatment in the I-R HT group restored HIF-1a expression to levels significantly higher than in the SHAM HT+PRL group (*p* < 0.05), indicating a rescuing effect of prolactin. In contrast, WT males treated with PRL (both SHAM and I-R) showed minimal HIF-1a expression, suggesting a suppressive effect of prolactin on this pathway in the wild-type background.

These findings highlight a complex regulatory pattern in which PRL modulates HIF-1a expression in a genotype- and sex-specific manner, suppressing activation in WT animals while rescuing or downregulating it in HT contexts, particularly following injury.

### VEGF Expression Indicates Prolactin-Induced Angiogenic Activation in a Sex- and Genotype-Dependent Manner

Vascular Endothelial Growth Factor (VEGF) expression was analyzed to assess angiogenic responses under different injury and treatment conditions. In male mice, VEGF levels were minimal in all non-PRL-treated groups (SHAM and I-R, WT and HT), suggesting that neither baseline nor IRI alone was sufficient to induce angiogenesis (Figure 7a). However, PRL-treated groups exhibited a marked increase in VEGF expression. The highest levels were observed in the I-R HT+PRL group, followed by I-R WT+PRL and SHAM HT+PRL. These results indicate that prolactin induces a strong pro-angiogenic response in male mice, particularly under injury and in heterozygous backgrounds.

**Figure 7.**
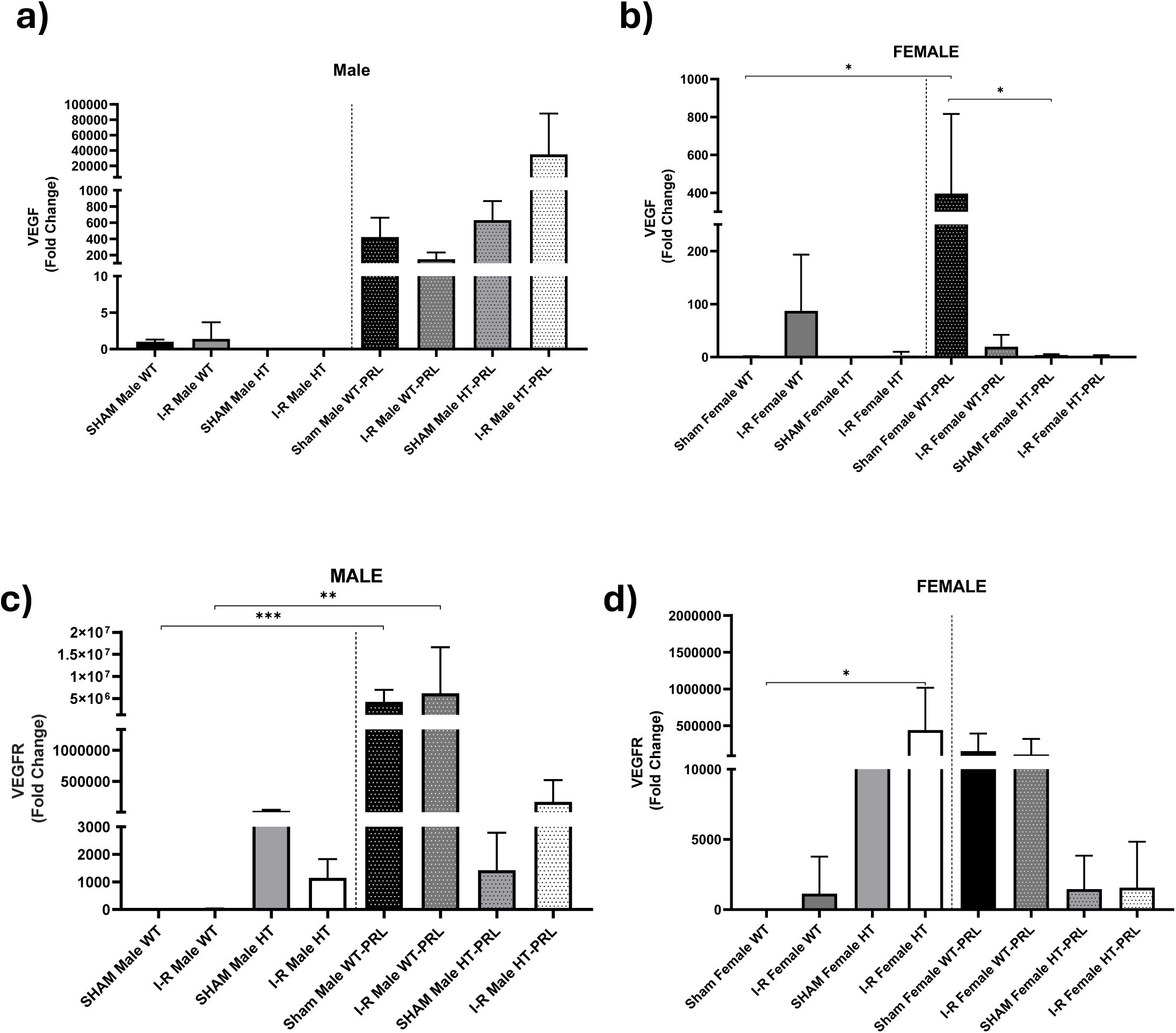
Prolactin induces VEGF and VEGFR expression in a sex- and genotype-dependent manner. Renal mRNA levels (RT-qPCR; fold-change normalized to Gapdh) of VEGF in (a) male and (b) female mice, and of its receptor VEGFR in (c) male and (d) female mice, across genotype, injury and treatment groups. (a) In males, VEGF was low in all untreated groups and strongly induced by PRL, with the highest levels in I-R HT+PRL. (b) In females, VEGF increased selectively in I-R WT+PRL (*p < 0.05 vs. I-R WT and sham HT+PRL), with negligible induction in HT females. (c) In males, VEGFR was markedly increased by PRL, peaking in the WT+PRL groups (***p < 0.001 and **p < 0.01 vs. sham WT and sham HT, respectively), with only a modest rise in I-R HT+PRL. (d) In females, sham HT showed higher basal VEGFR than sham WT (*p < 0.05), and PRL consistently reduced VEGFR, with the lowest levels in I-R HT+PRL. Data are mean ± SD; n ≥ 5 per group. *p < 0.05, **p < 0.01, ***p < 0.001.

In female mice, VEGF expression remained low in most conditions, with select exceptions (Figure 7b). The I-R WT+PRL group showed a significant increase in VEGF compared to both I-R WT and SHAM HT+PRL controls (*p* < 0.05), indicating that PRL enhances VEGF expression specifically in injured WT females. A similar upregulation was noted in SHAM WT+PRL compared to SHAM HT+PRL, suggesting that PRL-driven angiogenic signaling is more effective in the wild-type genotype. Conversely, all PRL-treated HT females displayed negligible VEGF expression regardless of injury status.

### VEGFR Expression Highlights Divergent Regulation by Prolactin Across Sex and Genotype

VEGFR (Vascular Endothelial Growth Factor Receptor) expression exhibited strong sex- and genotype-dependent variation in response to prolactin and ischemic injury. In male mice, PRL treatment markedly increased VEGFR levels, with the highest expression observed in Sham WT+PRL and I-R WT+PRL groups, followed closely by Sham HT+PRL (Figure 7c). These levels were significantly higher than those seen in Sham WT and Sham HT controls (\*\**p* < 0.001 and \**p* < 0.01, respectively), suggesting a robust prolactin-induced angiogenic priming in WT males, both at baseline and post-injury. I-R HT+PRL males showed only a modest increase, with VEGFR expression remaining significantly lower than WT+PRL groups. Minimal VEGFR levels were observed in I-R HT and Sham WT males.

In females, a distinct pattern emerged. Sham HT mice exhibited significantly higher VEGFR expression than Sham WT counterparts (*p* < 0.05), reflecting genotype-driven basal upregulation (Figure 7d). Following IRI, VEGFR levels remained moderately high in both WT and HT animals without major differences. However, prolactin treatment consistently reduced VEGFR expression in both genotypes. The I-R HT+PRL group demonstrated the lowest VEGFR levels across all female conditions, underscoring a suppressive effect of prolactin in the context of injury and heterozygosity.

### TGF-p Expression in Mice Under Ischemia-Reperfusion and Prolactin Treatment

TGF-β expression was markedly elevated in I-R Female WT and SHAM Female WT, with a significant difference between SHAM WT and SHAM HT (*p < 0.05), indicating a basal and injury-related activation of TGF-β in wild-type females (Figure 8a). This suggests an intrinsic susceptibility of WT females to TGF-β-mediated processes such as fibrosis or immune regulation.

**Figure 8.**
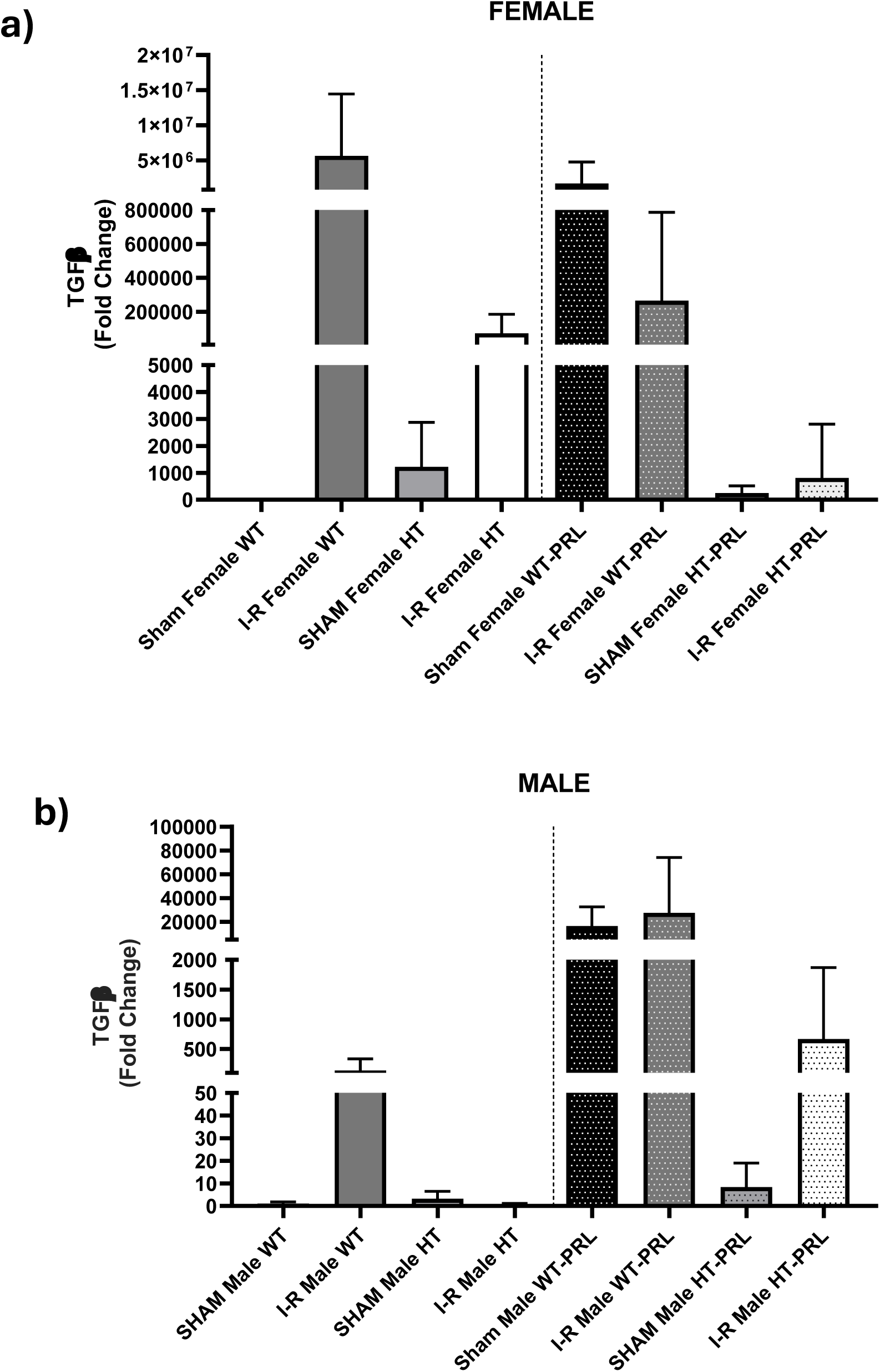
TGF-P expression reflects fibrotic potential and is modulated by prolactin. Renal TGF-β mRNA (RT-qPCR; fold-change normalized to Gapdh) in (a) female and (b) male mice. (a) In females, TGF-β was elevated in WT animals under both sham and I-R conditions, with a significant difference between sham WT and sham HT (*p < 0.05); PRL sustained high TGF-β in WT females, whereas HT+PRL groups showed markedly reduced expression. (b) In males, TGF-β was low in untreated groups and strongly increased by PRL in both genotypes, with the highest levels in WT+PRL animals. Note the segmented (broken) y-axis. Data are mean ± SD; n ≥ 5 per group. *p < 0.05.

Prolactin treatment maintained high TGF-β levels in WT females under both sham and I-R conditions, especially in the I-R Female WT+PRL group, suggesting that PRL sustains or enhances TGF-β signaling in injured WT females. In contrast, HT+PRL groups (sham and I-R) displayed markedly reduced TGF-β expression, highlighting a protective or repressive interaction between prolactin and the heterozygous genotype in modulating fibrotic responses.

Transforming Growth Factor Beta (TGF-β) expression remained low in non-prolactin-treated male groups (Figure 8b), with a modest increase in the I-R Male WT group, indicating a limited fibrotic or regulatory response to ischemic injury alone.

In contrast, prolactin treatment markedly increased TGF-β expression in both sham and I-R conditions, especially in WT and HT backgrounds. The I-R Male WT+PRL and SHAM Male WT+PRL groups exhibited the highest TGF-β levels, with similarly elevated values in HT+PRL conditions.

These results suggest that prolactin stimulates TGF-β expression in male kidneys, regardless of injury status or genotype, although WT mice tend to show a more pronounced response.

### IL-6 Expression in Male and Female Mice Under Ischemia-Reperfusion and Prolactin Treatment

In female mice, IL-6 expression was significantly upregulated in response to IRI, with a further increase observed in the group overexpressing PRL (Figure 9a). However, this effect was mitigated in the heterozygous PRL-overexpressing group, in which IL-6 levels were significantly reduced compared to the PRL-only group. These findings suggest that while PRL promotes IL-6–mediated inflammation following renal injury, Cathepsin D deficiency partially counteracts this proinflammatory effect.

**Figure 9.**
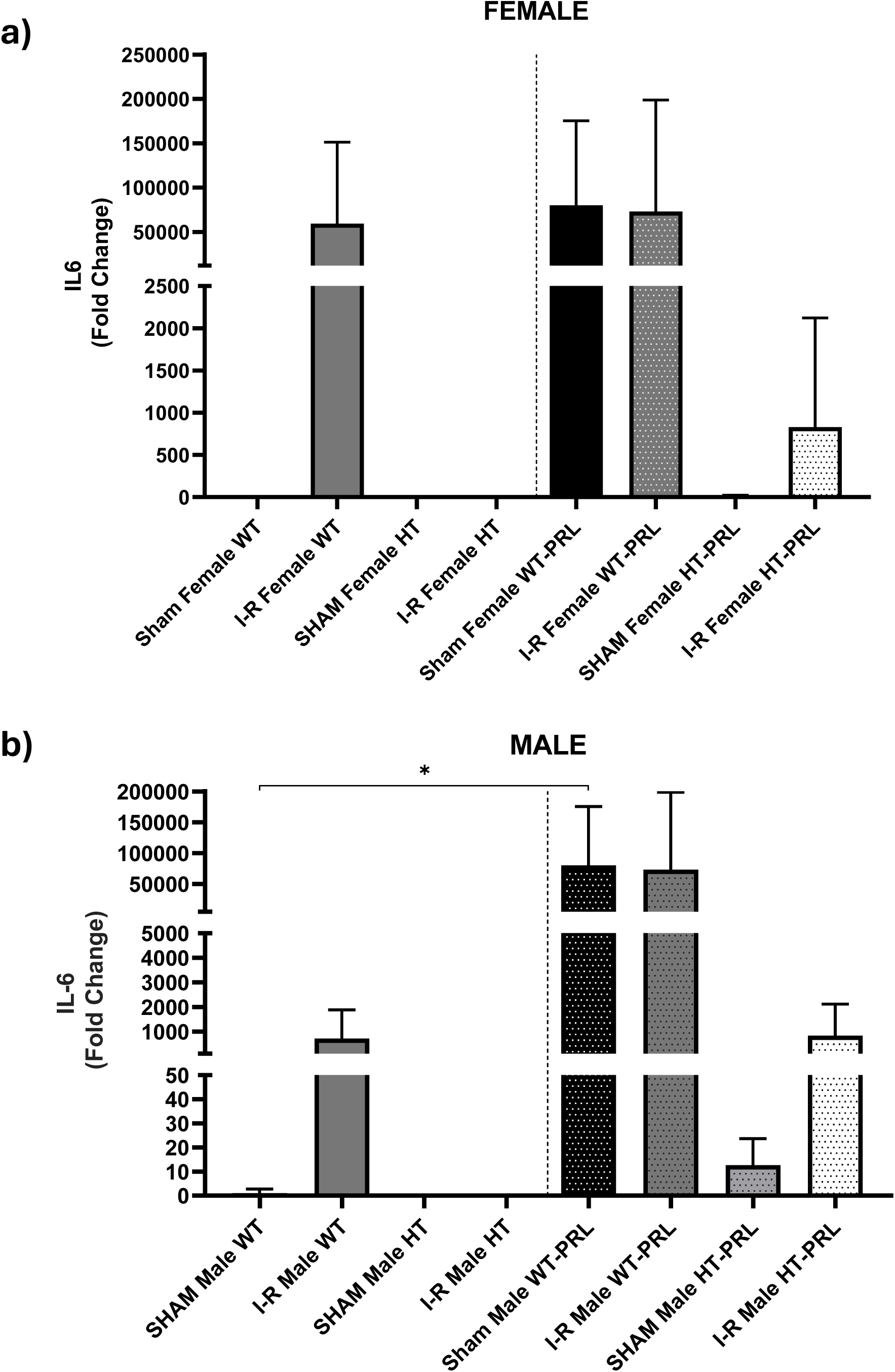
IL-6 expression is induced by ischemia–reperfusion and prolactin in a genotype-dependent manner. Renal IL-6 mRNA (RT-qPCR; fold-change normalized to Gapdh) in (a) female and (b) male mice. (a) In females, IL-6 was induced by I-R and further increased by PRL in the WT background, whereas HT+PRL animals showed markedly lower IL-6, indicating that Cathepsin D deficiency attenuates the PRL-driven proinflammatory response. (b) In males, IL-6 increased after I-R in WT animals and was strongly amplified by PRL in both genotypes (*p < 0.05 vs. sham WT), with comparatively lower induction in I-R HT+PRL. Note the segmented (broken) y-axis. Data are mean ± SD; n ≥ 5 per group. *p < 0.05.

IL-6 expression was significantly increased in the I-R Male WT group compared to the SHAM Male WT group (*p < 0.05), indicating a robust pro-inflammatory response triggered by ischemia-reperfusion injury (Figure 9b). Remarkably, PRL treatment in both WT and HT backgrounds resulted in a massive increase in IL-6 expression, reaching fold changes of several orders of magnitude higher than in untreated groups. This suggests that prolactin markedly enhances IL-6 transcription, regardless of the presence of injury.

Among prolactin-treated animals, the I-R Male HT+PRL group showed a relatively lower, but still elevated, IL-6 expression compared to PRL-treated WT animals, indicating a genotype-dependent modulation of the inflammatory response.

### Protein Identification Reveals Renal-Relevant Targets Modulated by Ischemia and Prolactin

Proteomic profiling of kidney samples extracted using RIPA buffer and analyzed via Progenesis QI identified a set of proteins previously reported in renal tissue (Table 1). Among the most confidently detected were Acyl-CoA:lysophosphatidylglycerol acyltransferase 1, Synaptophysin, and Ceramide-1-phosphate transfer protein, all involved in lipid processing, membrane remodeling, or vesicular trafficking relevant to renal physiology. Proteins such as SH3 domain-binding protein 5-like (Q99LH9) and Gasdermin-A (Q9EST1) were identified with high confidence scores and multiple unique peptides, suggesting strong and condition-dependent expression.

**Table 1.** Differentially expressed renal proteins identified by label-free LC-MS/MS (Progenesis QI). Selected high-confidence proteins detected in RIPA extracts of renal cortex and previously reported in kidney tissue. For each protein, the table lists the UniProt accession, peptide count, number of unique peptides, confidence score, ANOVA p-value, q-value (FDR-adjusted), molecular mass (Da), and description. Proteins were considered significant at q < 0.05.

| Accession | Peptide count | Unique peptides | Confidence score | Anova (p) | q Value | Mass | Description |
| --- | --- | --- | --- | --- | --- | --- | --- |
| Q91YX5 | 1 | 1 | 4.7644 | 0.01659107 | 0.0008763 | 43088.5843 | Acyl-CoA:lysophosphatidylglycerol acyltransferase 1 |
| Q62277 | 2 | 2 | 16.2633 | 0.07205689 | 0.00165553 | 34309.823 | Synaptophysin |
| Q6PGB6 | 1 | 1 | 4.9998 | 0.1675274 | 0.00165553 | 19414.4019 | N-alpha-acetyltransferase 50 |
| Q8BS40 | 1 | 1 | 4.2446 | 0.20423469 | 0.00165553 | 24597.2767 | Ceramide-1-phosphate transfer protein |
| Q8BFZ8 | 1 | 1 | 8.8256 | 0.23638653 | 0.00165553 | 16626.3645 | Protein FAM72A |
| Q8R115 | 2 | 2 | 9.5939 | 0.25623079 | 0.00165553 | 38881.8006 | Transmembrane protein 82 |
| O88890 | 1 | 1 | 3.8952 | 0.2586952 | 0.00165553 | 13903.8125 | SH2 domain-containing protein 1A |
| Q99LH9 | 4 | 4 | 15.4351 | 0.27668609 | 0.00165553 | 43374.141 | SH3 domain-binding protein 5-like |
| P09174 | 1 | 1 | 4.4783 | 0.31591769 | 0.00165553 | 9637.1893 | Retinal rod rhodopsin-sensitive cGMP 3'_5'-cyclic phosphodiesterase subunit-γ |
| Q8VCM5 | 5 | 5 | 18.5639 | 0.37075364 | 0.00165553 | 39835.0807 | Mitochondrial ubiquitin ligase activator of NFKB 1 |
| P32442 | 2 | 2 | 3.8476 | 0.37258094 | 0.00165553 | 27980.0116 | Homeobox protein MOX-1 |
| Q9EST1 | 1 | 1 | 12.2108 | 0.37613142 | 0.00165553 | 49593.1879 | Gasdermin-A |
| A8R0V4 | 2 | 2 | 20.3408 | 0.43846177 | 0.00176827 | 12569.9066 | Exocrine gland-secreted peptide 22 |
| Q8BY98 | 1 | 1 | 4.8769 | 0.46870423 | 0.00176827 | 36296.8992 | Ankyrin repeat domain-containing protein SOWAHD |
| Q3UTS8 | 2 | 2 | 4.5333 | 0.56397585 | 0.00198585 | 11530.4834 | Serine protease inhibitor Kazal-type 13 |
| Q9Z262 | 1 | 1 | 4.4252 | 0.64703644 | 0.00213593 | 23387.8162 | Claudin-6 |

Additional hits included Mitochondrial ubiquitin ligase activator of NF-κB 1 (Q8VCM5) and N-alpha-acetyltransferase 50 (Q6PGB6), indicative of mitochondrial stress response and post-translational modification pathways. These findings align with the inflammatory and fibrotic signatures identified in transcriptomic and cytokine profiling. The fact that all proteins have previously been validated in renal tissue underscores the robustness and biological relevance of the proteomic approach employed.

To gain mechanistic insights into the molecular responses associated with IRI and prolactin modulation, we conducted a pathway enrichment analysis on the differential expression data (Figure 10). The results revealed prominent enrichment in pathways involved in inflammation, cellular metabolism, and signal transduction.

**Figure 10.**
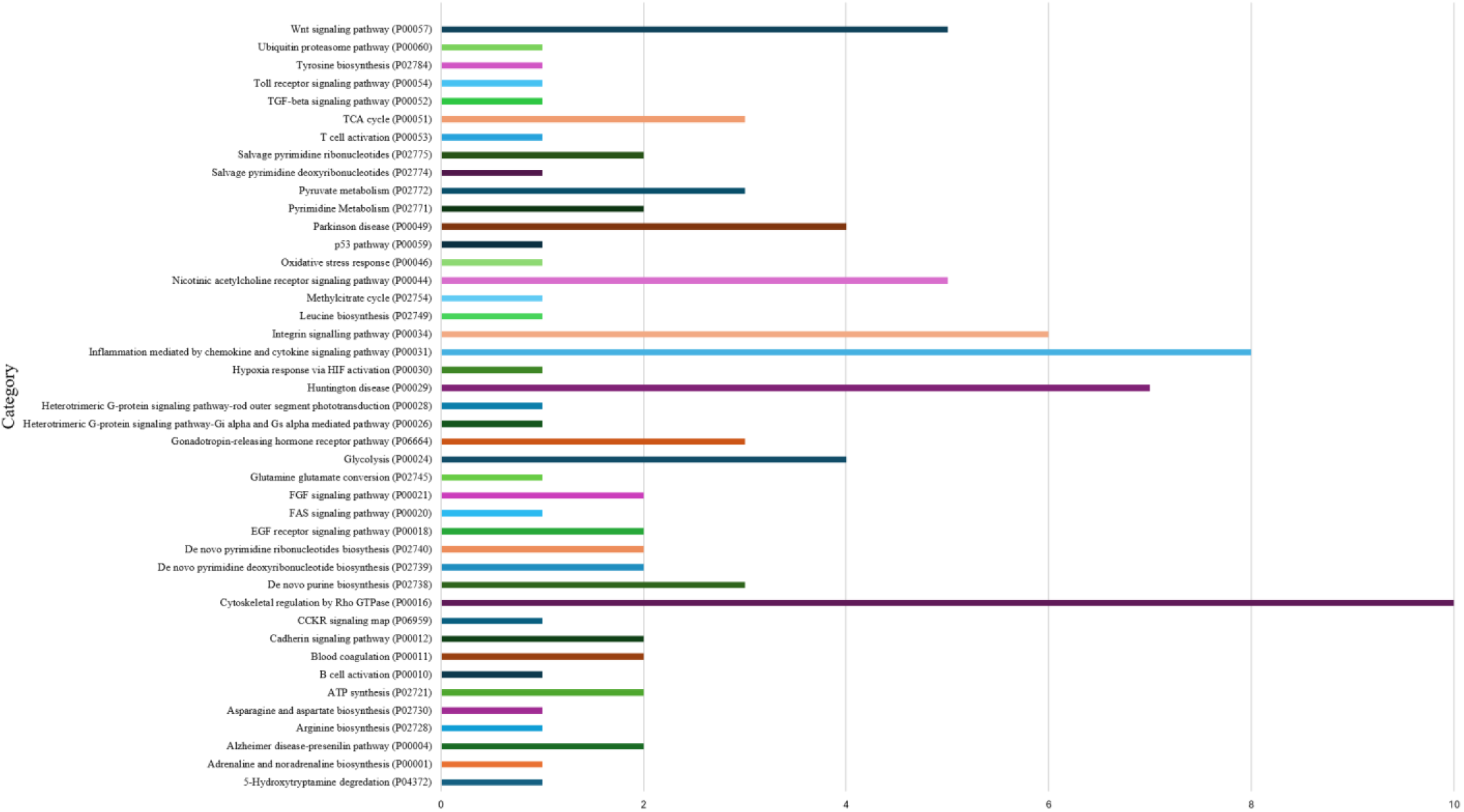
Pathway enrichment of proteins modulated by ischemia–reperfusion and prolactin. Functional enrichment of differentially expressed renal proteins, displayed as a horizontal bar plot of enriched pathways (PANTHER pathway identifiers in parentheses). Enriched categories included inflammatory signaling, metabolic remodeling, and additional signal-transduction pathways. The x-axis represents the enrichment metric. Pathways were considered significantly enriched at p < 0.05 (Benjamini–Hochberg FDR).

Among the most frequently enriched categories were the CCR4 signaling pathway (P05939), Huntington’s disease (P00029), and inflammation mediated by chemokine and cytokine signaling (P00031). Notably, inflammatory signaling pathways, including hypoxia response via HIF activation (P00030) and TGF-β signaling (P00052), were consistently enriched, consistent with the pro-inflammatory and pro-fibrotic profile observed in our expression analyses.

Metabolic pathways such as pyruvate metabolism (P02772), glycolysis (P00024), and amino acid biosynthesis and degradation (e.g., leucine and tyrosine biosynthesis, P02749 and P02740) were also represented, suggesting active metabolic remodeling post-injury.

Furthermore, several neurotransmitter-related pathways—including nicotinic acetylcholine receptor signaling (P00044) and 5-hydroxytryptamine degradation (P04372)—were enriched, possibly reflecting neuroimmune crosstalk in renal stress contexts.

Overall, these findings highlight a complex interplay between immune activation, hypoxic stress adaptation, and hormone-regulated signaling, consistent with the observed modulation of HIF-1, IL-6, VEGF, and TGF-β expression across experimental groups.

## Discussion

This study suggests a sex-dependent role of Cathepsin D and prolactin signaling in modulating the renal response to IRI. Our findings reveal that Cathepsin D is significantly upregulated in response to IRI, particularly in female mice, and that it functions enzymatically to cleave PRL into Vasoinhibins under physiological conditions. These Vasoinhibins fragments are emerging as critical effectors in renal physiology, particularly in genetically susceptible backgrounds.

Our data aligned with previous reports indicating that PRL has both immunomodulatory and tissue protective functions. Several studies have shown that PRL can reduce inflammation and oxidative stress in cardiovascular and renal models of injury. For instance, PRL has been implicated in promoting endothelial repair and angiogenesis via VEGF-dependent pathways [24, 25]. These observations are consistent with our findings that PRL upregulated VEGF and VEGFR expression in WT males, and to a lesser degree in HT animals. However, contrasting data also exist in some contexts, prolactin has been associated with fibrotic responses, particularly in models of diabetic nephropathy or autoimmune disease, where PRL signaling promotes TGF-β and extracellular matrix deposition [26, 27]. In our study, PRL enhanced TGF-β expression in WT animals but not in HT mice, suggesting a genotype-dependent divergence that may reconcile these conflicting observations.

Sex differences in the renal response to injury are well-documented, with females generally exhibiting lower susceptibility to IRI, attributed in part to estrogen-mediated signaling and mitochondrial resilience [8]. Our findings echo these patterns, as WT females showed moderate injury post-IRI and improved significantly with PRL treatment, while HT females appeared intrinsically protected. In contrast, HT males were more vulnerable to IRI and benefitted markedly from PRL, a response not seen in WT males. These sex-specific findings highlight the complexity of hormonal-immune interactions and underscore the need for sex-balanced research in nephrology.

Another key contribution of this study is the identification of Cathepsin D as a mediator of PRL cleavage, extending previous in vitro work [28]. This enzymatic axis provides a mechanistic link between lysosomal protease activity and endocrine regulation of inflammation. However, the biological relevance of Vasoinhibins remains controversial. While their antiangiogenic properties have been well described in ocular and tumor biology, their role in renal physiology is less understood and merits further investigation. Proteomic and pathway enrichment analyses revealed consistent activation of metabolic stress and inflammatory signaling in IRI models, including CCR4 signaling, glycolysis, and oxidative stress pathways. Interestingly, our PRL-treated groups showed modulation of these signatures, suggesting a reprogramming of injury response. Some of these effects contradict earlier studies that described PRL as pro-inflammatory in autoimmune contexts [29], underscoring the tissue- and context-specific actions of PRL.

Together, these findings highlight a complex interplay between prolactin, Cathepsin D, sex, and genetic background in modulating kidney injury and repair. Our study suggests that PRL’s renoprotective actions are amplified in genetically sensitized contexts (HT) and sex-specific milieus, offering mechanistic insight and potential therapeutic directions. Nonetheless, future work is needed to dissect the full repertoire of Vasoinhibins effects in the kidney, explore the long-term consequences of PRL modulation, and evaluate translational potential in human models of acute kidney injury.

## Conclusion

Our findings underscore the critical role of Cathepsin D and prolactin in mediating sex- and genotype-specific responses to renal ischemia-reperfusion injury. We demonstrate that Cathepsin D not only responds robustly to renal injury, but also functions enzymatically to generate Vasoinhibins from prolactin under physiological conditions. These fragments, in turn, influence key pathways involved in inflammation, angiogenesis, fibrosis, and renal function preservation.

Prolactin treatment exerted protective effects in both male and female mice, but its efficacy was highly dependent on genetic background, with heterozygous animals exhibiting more favorable modulation of pro-inflammatory and pro-fibrotic markers. These results suggest that partial Cathepsin D deficiency may create a regulatory environment conducive to PRL’s beneficial effects, in part by altering the balance between PRL and Vasoinhibins activity.

This work highlights the therapeutic potential of targeting PRL and Cathepsin D in acute kidney injury and suggests that consideration of patient sex and genetic background may be critical in designing future precision therapies. Further studies are warranted to evaluate the long-term impact of this hormonal regulation in chronic kidney disease models and to validate translational applicability in human populations.

---

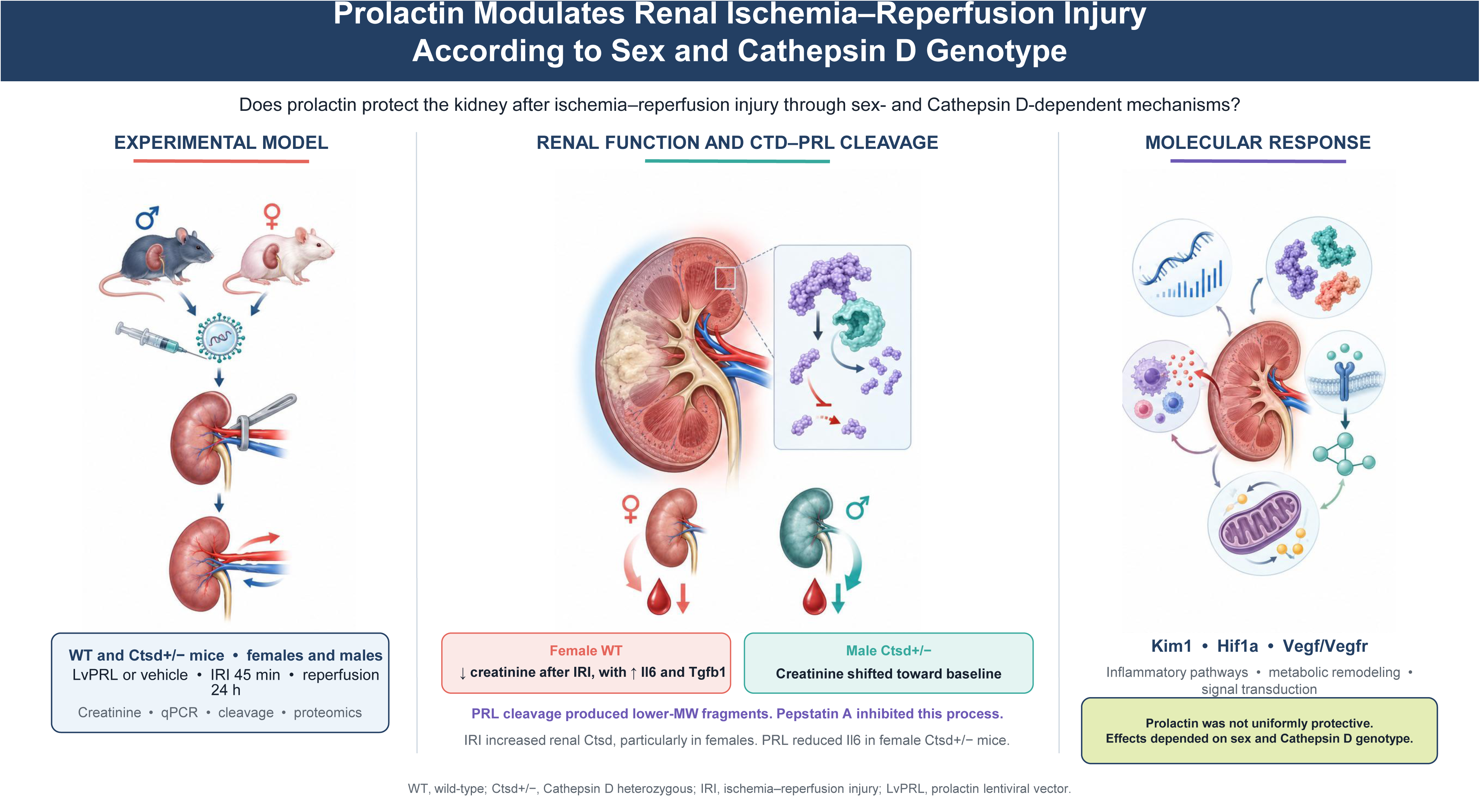

## References

1. Kadatane, S.P., et al. The Role of Inflammation in CKD. Cells, 2023.12(12).

2. Rayego-Mateos, S., et al., Molecular Mechanisms of Kidney Injury and Repair. Int J Mol Sci, 2022. 23(3).

3. Ciarambino, T., P. Crispino, and M. Giordano, Gender and Renal Insufficiency: Opportunities for Their Therapeutic Management? Cells, 2022. 11(23).

4. Kang, K.P., et al., Effect of gender differences on the regulation of renal ischemia-reperfusion-induced inflammation in mice. Mol Med Rep, 2014. 9(6): p. 2061–8.

5. Valdivielso, J.M., C. Jacobs-Cacha, and MJ. Soler, Sex hormones and their influence on chronic kidney disease. Curr Opin Nephrol Hypertens, 2019. 28(1): p. 1–9.

6. Franco-Acevedo, A., R. Echavarria, and Z. Melo, Sex Differences in Renal Function: Participation of Gonadal Hormones and Prolactin. Endocrines, 2021. 2(3): p. 185–202.

7. Lima-Posada, I. and N.A. Bobadilla, Understanding the opposite effects of sex hormones in mediating renal injury. Nephrology (Carlton), 2021. 26(3): p. 217–226.

8. Lima-Posada, I., et al., Gender Differences in the Acute Kidney Injury to Chronic Kidney Disease Transition. Sci Rep, 2017. 7(1): p. 12270.

9. Saleem, M., H. Martin, and P. Coates, Prolactin Biology and Laboratory Measurement: An Update on Physiology and Current Analytical Issues. Clin Biochem Rev, 2018. 39(1): p. 3–16.

10. Rojas-Vega, L., et al., Ovarian hormones and prolactin increase renal NaCl cotransporter phosphorylation. Am J Physiol Renal Physiol, 2015. 308(8): p. F799–808.

11. Dourado, M., et al, Relationship between Prolactin, Chronic Kidney Disease, and Cardiovascular Risk. Int J Endocrinol, 2020. 2020: p. 9524839.

12. Huang, S., et al., Research progress on the role of hormones in ischemic stroke. Front Immunol, 2022. 13: p. 1062977.

13. Clapp, C., S. Thebault, and G. Martinez de la Escalera, Role of prolactin and vasoinhibins in the regulation of vascular function in mammary gland. J Mammary Gland Biol Neoplasia, 2008. 13(1): p. 55–67.

14. Moreno-Carranza, B., et al, Prolactin promotes normal liver growth, survival, and regeneration in rodents: effects on hepatic IL-6, suppressor of cytokine signaling-3, and angiogenesis. Am J Physiol Regul Integr Comp Physiol, 2013. 305(7): p. R720–6.

15. Hilfiker-Kleiner, D., et al, A cathepsin D-cleaved 16 kDa form of prolactin mediates postpartum cardiomyopathy. Cell, 2007. 128(3): p. 589–600.

16. Morohoshi, K., et al., 16 kDa vasoinhibin binds to integrin alpha5 beta1 on endothelial cells to induce apoptosis. Endocr Connect, 2018. 7(5): p. 630–636.

17. Robles, J.P., et al., Vasoinhibin’s Apoptotic, Inflammatory, and Fibrinolytic Actions Are in a Motif Different From Its Antiangiogenic HGR Motif. Endocrinology, 2023.165(2).

18. Triebel, J., et al., Principles of the prolactin/vasoinhibin axis. Am J Physiol Regul Integr Comp Physiol, 2015. 309(10): p. R1193–203.

19. Yamamoto-Nonaka, K., et al, Cathepsin D in Podocytes Is Important in the Pathogenesis of Proteinuria and CKD. J Am Soc Nephrol, 2016. 27(9): p. 2685–700.

20. Cocchiaro, P., et al., The Multifaceted Role of the Lysosomal Protease Cathepsins in Kidney Disease. Front Cell Dev Biol, 2017. 5: p. 114.

21. Ozkayar, N., et al., Relation between serum cathepsin D levels and endothelial dysfunction in patients with chronic kidney disease. Nefrologia, 2015. 35(1): p. 72–9.

22. Radcliffe, P.A., et al., Analysis of factor VIII mediated suppression of lentiviral vector titres. Gene Ther, 2008. 15(4): p. 289–97.

23. Adan, N., et al.. Prolactin promotes cartilage survival and attenuates inflammation in inflammatory arthritis. J Clin Invest, 2013.123(9): p. 3902–13.

24. Goldhar, A.S., et al., Prolactin-induced expression of vascular endothelial growth factor via Egr-1. Mol Cell Endocrinol, 2005. 232(1-2): p. 9–19.

25. Zhao, H., et al., The role of prolactin/vasoinhibins in cardiovascular diseases. Animal Model Exp Med, 2023.6(2): p. 81–91.

26. Li, Y., et al., Correlation between serum prolactin and the systemic immune-inflammation index in diabetic kidney disease: a cross-sectional study. Front Endocrinol (Lausanne), 2026. 17: p. 1772810.

27. Borba, V.V., G. Zandman-Goddard, and Y. Shoenfeld, Prolactin and Autoimmunity. Front Immunol, 2018. 9: p. 73.

28. Cruz-Soto, M.E., et al., Cathepsin D is the primary protease for the generation of adenohypophyseal vasoinhibins: cleavage occurs within the prolactin secretory granules. Endocrinology, 2009. 150(12): p. 5446–54.

29. Brand, J.M., et al., Prolactin triggers pro-inflammatory immune responses in peripheral immune cells. Eur Cytokine Netw, 2004. 15(2): p. 99–104.

